# Sex Differences in Plasma Levels of Endocannabinoids and Related Lipids Before and After acute and repeated mTBI: an exploratory study for plasma biomarkers for mTBI

**DOI:** 10.1101/2025.09.10.675401

**Authors:** Emily Richter, Taylor Woodward, Praveen P Kulkarni, Craig F. Ferris, Heather B Bradshaw

## Abstract

Mild traumatic brain injury (mTBI) is common diagnosis across all age groups and while most symptoms resolve within a few weeks; between 10 and 25 percent of mTBI patients suffer long-term problems. Known as post-concussion syndrome (PCS), symptoms include headache, a range of cognitive deficits, and depression. Currently, there are no established treatments for PCS and no clear predictive biometrics to determine which patients are at increased risk. Previous studies have identified some protein-derived plasma biomarkers for mTBI, however, the effects of mTBI on lipid signaling molecules and metabolites in blood is largely unknown. Endogenous lipids (endolipids) such as the endocannabinoids (eCBs) and their congeners are lipid signaling molecules that are associated with promoting neuroprotective responses after head trauma in animal models. Here, we examine the plasma lipidome using a rat model of acute and repeated mTBI that we previously demonstrated had a sex dependent change in neuroinflammation wherein females showed a higher degree of neurodegeneration after repeated head-injury than males. Key results of this exploratory lipidomics screen here demonstrates that acute head injury drives significantly more changes in plasma endolipids in males (32%) than females (8%), whereas, on the second day of head injury, only 11% change in males but 15% in females. Some key endolipids were modified in both males are precursors for resolving molecules and this was lacking in females. Given that females with repeated mTBI in this model demonstrated aspects of PCS, this could be an important component in evaluating clinical cases. Endolipids in the screen were measurable in plasma using only 100µL, a volume necessary to be able to perform multiple blood draws on these rodent subjects. This threshold provides evidence that the levels of these endolipids could be readily measured throughout a patient’s recovery. Therefore, this family of endolipids has the potential to provide data on the progression of the injury and could be another crucial aspect in predicting mTBI outcomes.

## Introduction

Mild traumatic brain injury (mTBI) is an extremely common diagnosis across all age groups from pediatric to geriatric and can result from a multitude of events, including sports, work, and everyday life. In 2020 there were approximately 586 hospitalizations per day due to head-related injuries [1], not including the number which go untreated. About 70-90% of these injuries would qualify as mild brain injuries [2] and can be resolved within a few weeks; however, between 10 and 25 percent of mTBI patients suffer long-term problems. Known as post-concussion syndrome (PCS), symptoms are predominately headache, a range of cognitive deficits, and depression [1]. Currently, there are no established treatments for PCS and no clear predictive biometrics to determine which patients are at increased risk [2]. Previous studies have investigated the effects of mTBI on certain time-dependent proteins in both animal and clinical models, for hours and months after injury [2-8]. However, the effects of mTBI on lipid signaling molecules and metabolites in blood is largely unknown.

Endogenous cannabinoids, or endocannabinoids (eCBs) are lipid signaling molecules that are associated with promoting neuroprotective responses after head trauma in animal models [9]. The canonical eCBs, *N*-arachidonoyl ethanolamine (AEA; Anandamide) and 2-arachidonyl glycerol (2AG) are associated with modulating inflammatory responses and excitotoxicity prevention [10]. eCBs modulate responses throughout the nervous system and in the periphery, at least in part, via the cannabinoid CB1 and CB2 receptors signaling [9, 10], however, these effects are not limited to these signaling systems. These eCBs are only two of over 100 endogenous congeners that the Bradshaw lab and others have shown to share biosynthetic and metabolic pathways [11-17], are modulated by environmental toxins [18], and drugs of abuse [15, 19, 20]. We have also demonstrated changes in these lipids in a rat model of mTBI focusing primarily on the trigeminal nucleus, which was the most effected after a single cranial hit to simulate mTBI, with significant increases in the concentration of the many of the endolipids we evaluate, including the eCBs and their congeners [21]. Each of these prior studies evaluated

these classes of lipid signaling molecules in CNS tissue, however, there is a need to understand any effects on circulating endolipids in the bloodstream, which can add to the knowledge of potential biomarkers in both rodents and humans. We have demonstrated the use of plasma in both human [22, 23] and animal studies [24] to determine changes in this large class of signaling lipids providing a basis for evaluating plasma lipids as potential biomarkers in mTBI.

An emerging literature on mTBI therapeutics and outcomes focuses on sex differences. Recent reviews on the topic provide leading hypotheses for outcome differences being that females may exhibit some types of neuroprotection associated with estrogenic hormones, though mechanisms of these sex differences are likely more complex and focused on acute measurements [25]. Previous research from the Ferris lab demonstrated that repeated mTBI in male and female rats resulted in significantly greater anisotropy (an MRI measure of diffusion coefficient that relates to tissue damage) in females at 12 days post injury verses 6 days post injury, suggesting that some of the initial parameters that were interpreted as protection (*e*.*g*. lower inflammatory mediators, BDNF, etc.) may result in longer term protection for males [26, 27]. Therefore, here we test the hypothesis that there are sex differences in plasma endolipid regulation in an mTBI model and that this is dependent on acute or repeated head trauma. These studies represent an exploratory study to identify potential biological markers of acute and successive head injury within and between sexes.

## Methods

### Subjects

Twenty adult male and female Sprague Dawley rats (n =10/group), 2-3 months old, weighing between 250–400g underwent two days (Day 1, D1 and Day 2, D2) of Head Hits (HH, n=10, n=5/per genetic sex), to produce mTBI, performed by a closed-head momentum exchange model. Data was collected during the light phase of the light/dark cycle. Subjects were obtained and maintained as described previously [28]. All methods and procedures described below were pre-approved by the Northeastern University Institutional Animal Care and Use Committee under protocol number 24-0517R-A1: MRI following Mild Head Injury. Northeastern University’s animal care and use program and housing facilities are fully accredited by AAALAC International. The protocols used in this study followed the ARRIVE guidelines for reporting *in vivo* experiments in animal research [29].

### mTBI momentum exchange model procedure

The momentum exchange model that was used to produce a head hit has been previously detailed in [30]. A pneumatic pressure drive 50g compactor [31] was used to reliably produce mild to severe head injuries to rats and then refined to test the behavioral effects of mild TBI controlling for the axis of injury, rotational force, and head acceleration [32]. Subjects were allowed to acclimate to animal handling 2 weeks before the head or shoulder hit. This momentum exchange model used an impact velocity of 7.4 m/s, which was determined by using high-speed video recordings. For head hits, the animals were restrained, and the impact piston was directed to the top of the skull, midline, in the approximate area of Bregma. The shoulder injury used the same restraint and apparatus, but with a novel protocol: the piston moved caudally and laterally along the midsagittal plane so that the impact position is perpendicular to the right scapula of the animal. The selection of the impact region was characterized by a large body of data on the effects of acute and long-term mTBI in rodents [33-36]. Hits were done under 2% isoflurane anesthesia, with a 24-hour interval between each impact. Rats were returned to their cages after the final mTBI or shoulder injury. All blood draws via tail vein and tests occurred during in the light phase of a normal light-dark cycle.

Whole blood samples were collected before the first head hit on each day to establish a baseline and then 15 minutes after a hit. Samples were centrifuged for 5 minutes, and the plasma was removed and stored at -80°C until shipped on dry ice to Indiana University Bloomington and was returned to -80°C until further processing. See Figure 1. for a visual representation of the experimental procedure.

**Figure 1.**
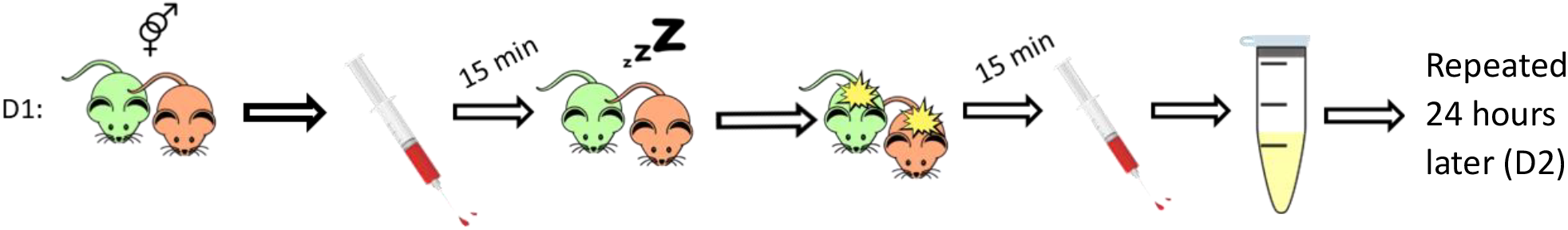
Visual Representation of Experimental Procedure: On Day 1 (D1), Rats were removed from their cages during the light phase of their light-dark cycle. Fifteen minutes prior to being anesthetized and subsequently receiving impact, whole blood was drawn from their tail veins. Once they were anesthetized, they were placed in the momentum exchange apparatus and received a single heat hit. After 15 minutes, they once again had their whole blood drawn from the tail vein. The blood was then centrifuged for 5 minutes, and the plasma was removed and stored at -80°C. This process was repeated after 24 hours for Day 2 (D2).

## Lipid Extraction and Partial Purification

Lipidomics analysis of plasma was performed as previously described [22, 24]. In brief, plasma samples (75µL) were vortexed and added to 2mL of HPLC-grade methanol (MeOH) + 5µL of 1µM 500 picomolar (pM) deuterium-labeled anandamide (d8AEA) as an internal standard to determine extraction efficiency. Samples were incubated in the dark on ice for 30 min before centrifugation (19,000 x g for 20 minutes at 20 °C). The supernatant was added to 8mL of HPLC-grade water for an approximately 25% organic solution and partially purified with Agilent C-18 solid phase extraction (SPE) columns (Santa Clara, CA, USA) as previously described. The following 1.0 mL elutions of 65-,75-, and 100-percent methanol were collected for High-Performance Lipid Chromatography/Mass Spec/Mass Spec (HPLC/MS/MS) and stored in a -80°C freezer until further. Samples were de-identified and recorded by processing day.

## HPLC/MS/MS analysis

HPLC/MS/MS analysis was performed as previously described [19]. In brief, samples were removed from the −80°C freezer and allowed to warm to room temperature and then vortexed for approximately 1 minute before being placed into the autosampler held at 24°C (Agilent 1100 series autosampler, Palo Alto, CA) for LC/MS/MS analysis. A 20µL injection volume from each sample was injected into a C18 Analytical Column (Agilent Technologies, Santa Clara, CA) to scan for compounds. Gradient elution (200 *μ*L/min) then occurred under the pressure created by two Shimadzu 10AdVP pumps (Columbia, MD) (mobile phase A: 20% HPLC methanol, 80% HPLC water, and 1 mM ammonium acetate; mobile phase B: 100% HPLC methanol and 1 mM ammonium acetate). Next, electrospray ionization was accomplished using an Applied Biosystems/MDS Sciex (Foster City, CA) API3000 triple quadrupole mass spectrometer. A

multiple reaction monitoring (MRM) setting on the LC/MS/MS was then used to analyze levels of each compound present in the sample injection. 16 lipid families were screened for a total of 86 individual compounds, one of which was the internal standard d8AEA. Synthetic standards were used to generate optimized MRM methods and standard curves for analysis [14, 16, 19].

## Statistical analysis

Statistical analyses were completed in IBM SPSS Statistics 29 (Chicago, IL,USA) as previously described [16]. Analysis of individual endogenous lipids in plasma in the following comparison groups were analyzed using Students t-tests set to 2-tails and Type 2: Male vs female baseline day 1, M vs F post hit D1; M vs F B Day 2, M vs F PH D2; F B vs F PH D1, M B vs M PH D1, F B vs F PH D2, M B vs M PH D2. Statistical significance for all tests was set at p < 0.05, and trending significance at 0.05 < p < 0.10. Descriptive and inferential statistics were used to create heatmaps for visualizing changes in the concentration of each lipid analyte for every condition. Briefly, the direction of changes for each analysis group compared to are depicted by color, with green representing an increase and orange representing a decrease. Level of significance is shown by color shade, wherein p < 0.05 is a dark shade and 0.05 < p < 0.1 is a light shade. Direction of the change compared to vehicle is represented by up (increase) or down (decrease) arrows. Effect size is represented by the number of arrows, where 1 arrow corresponds to 1–1.49-fold difference, 2 arrows to a 1.5–1.99-fold difference, 3 arrows to a 2– 2.99-fold difference, 4 arrows a 3–9.99-fold difference, and 5 arrows a difference of tenfold or more [37]. An abbreviation of ‘BDL’ indicates that the lipid concentration that was present in the sample was below the detectable levels of our equipment while ‘BAL’ indicates below analytical levels. To calculate the effect size of a lipid that was significantly higher in a particular group (male and female by baseline or post-hit by day for sex difference analysis and baseline and post-hit conditions by day of the same sex for the genetic sex analysis), the average concentration of the respective comparison group is divided by the average concentration of the vehicle group. To calculate the effect size of a compound that is significantly lower in the treatment group, the same calculation is conducted, except the inverse of the resulting ratio is used to represent a fold decrease. Finally, bar graphs which show data as mean ± s.e. mean were made using GraphPad Prism Software (La Jolla, CA, USA) for key lipid families.

## Results

### Overall effects of differences between and within sex and treatment groups on plasma lipids

Between the two days of head hits and subject groups (female: F and male: M), 86 endolipids in the screening library were detected in at least one plasma sample (*see Supplemental Figures 1 and 2 full list of endolipids screened and p values for all interactions analyzed*). Of these 86 compounds, 87.2% (75/86) met analytical limits for an exploratory study (*i*.*e*. detected in at least 3 out of 5 subjects). Of those 75, 69% (52/75) demonstrated significant differences in at least one comparison across all potential interactions.

Figure 3 provides a summary to the percentage and direction of change between subject and treatment groups. These are organized as sex differences at each of the four, blood collection time points (3A), then as treatment effects within genetic sex subject groups, then finally as combined (F and M) for each of the 2 days of head-hit treatments (3B). A summary of the sex difference analysis shows that levels of 31% of plasma endolipids at baseline on day 1 were significantly difference with 96% of those lipids being higher in females. The percentage of endolipids showing significant sex differences after the first head hit dropped to 14% and the direct was reversed with 82% being lower in females. Sex differences at baseline on day 2 of testing dropped to 9% and then 8% after head hit. In both cases the levels were lower in females 89% and 79% respectively.

**Figure 2.**
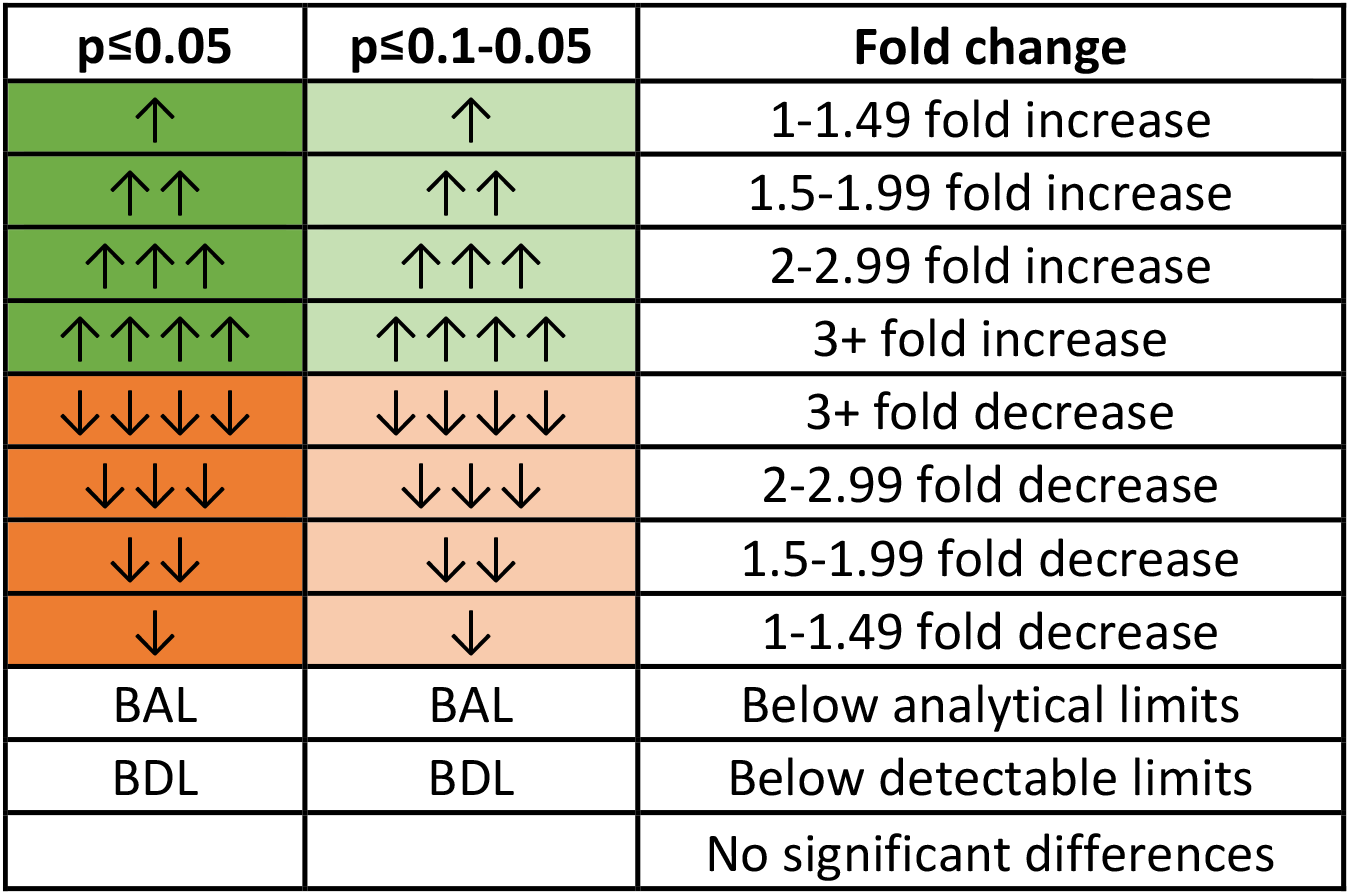
Heatmap Legend. Heatmaps are used to summarize means data and statistical significance to provide a shorthand for large-scale lipidomics datasets. Dark green represents either significant increases (after treatment) or levels that are significantly higher (sex differences) p ≤0.05 while light green represents significance levels of p ≤ 0.1-0.05. Dark orange represents significant decreases (after treatment) or levels that are significantly lower (sex differences) wherein p ≤0.05 while light orange represents significance levels of p≤ 0.1-0.05.Fold change is indicate d by the number of arrows, where 1 arrow corresponds to 1–1.49-fold difference, 2 arrows to a 1.5–1.99-fold difference, 3 arrows to a 2–2.99-fold difference, 4 arrows a 3–9.99-fold difference, and 5 arrows a difference of tenfold or more. BAL refers to endolipids that were measured in at least one sample but less than 4 making statistical analysis unreliable. BDL refers to endolipids that were not detected in any samples. Blank cells indicate that endolipids were present in at least 4 samples in each treatment group being analyzed and that no significant differences were present.

**Figure 3:**
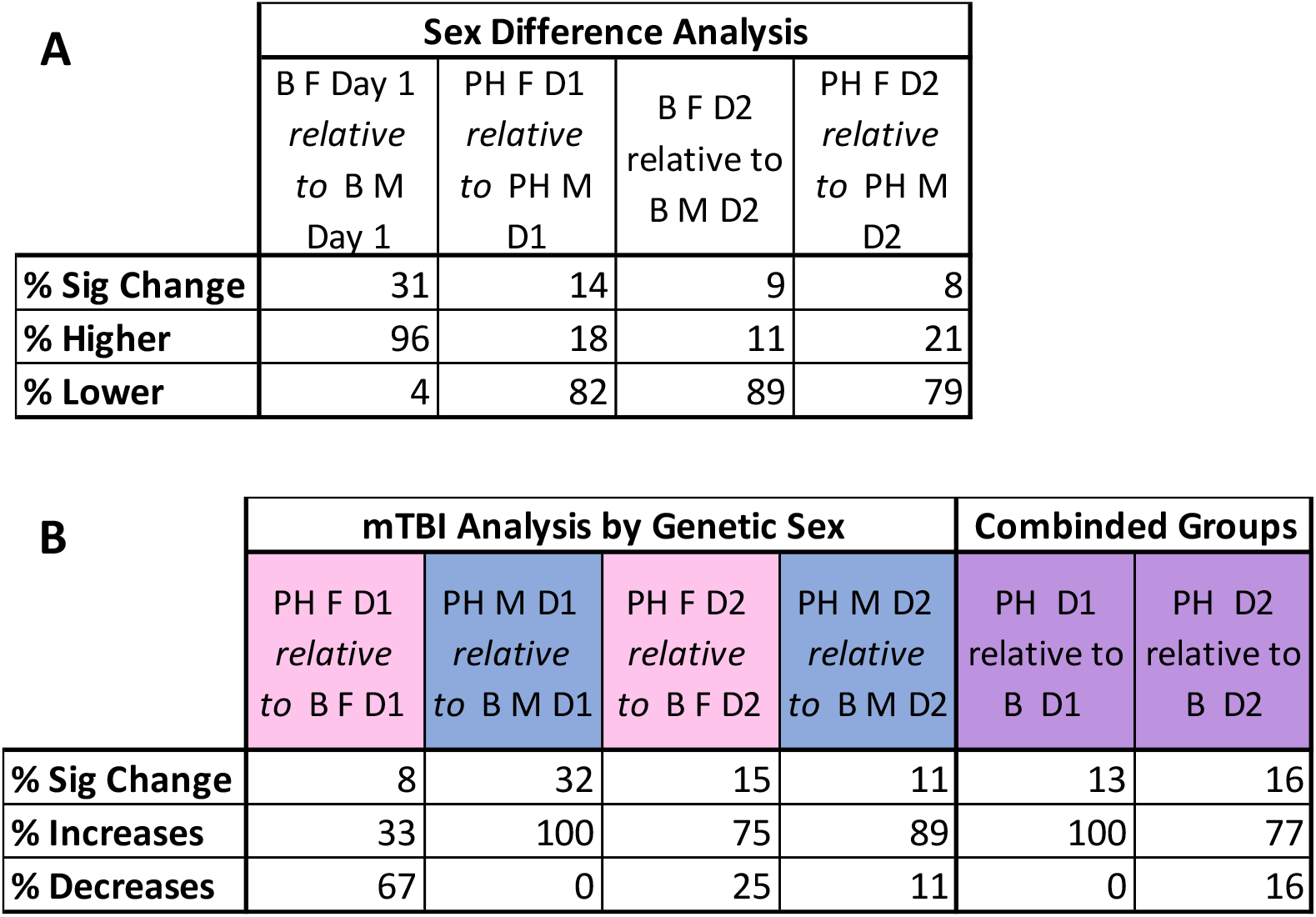
Summary of percent change in endolipids within and between groups. 3A shows percent changes (overall significant, higher, or lower) between females and males at each plasma evaluation timepoint in the study. 3B shows percent changes (overall significant, increases or decreases to treatment) within genetic sex and combined females and males comparing pre and post head hit.

The summary of acute and repeated heat hit effects by genetic sex (Fig. 3B) revealed that on day 1 of testing acute head hit caused only 8% of plasma lipids to change in females but 32% in males. Where the majority (67%) of these changes in females were decreases, 100% of the changes in males were increases. On day 2 of testing females showed 15% difference in plasma lipids after the second head-hit (75% increases) in 24 hours and males 11% (89% increases). Combining the female and male subjects provided a different outcome for day 1 data with a total of 13% of lipids changed (closer to the female percentage) after day 1 heat hit and 100% being increases (aligning with the male data). Day 2 combined results were more closing aligned with both female and male data with 16% overall change and 77% of that being increases.

### Lipidomics evaluations by subject and treatment group

Figures 4-6 illustrate the levels of 6 different endogenous lipids in each subject and treatment type point and then with the female and male data combined pre-post head hit on consecutive days. These examples include the endocannabinoids, Anandamide (4A) and 2-AG (4B), and their endogenous congeners stearoyl ethanolamine (5A), palmitoyl taurine (5B), linoleoyl glycine (6A) and oleoyl methionine (6B). For each example the mean with SEM is shown on the left. On the right a table shows the analytical interactions that are used to generate the larger heatmaps including fold-change difference, p value, and heatmap icon between the listed interaction (*e*.*g*. Baseline Female Day 1: BF D1 versus Baseline Male Day 1: BM D1; *Supplemental Figures for all analyses*).

**Figure 4.**
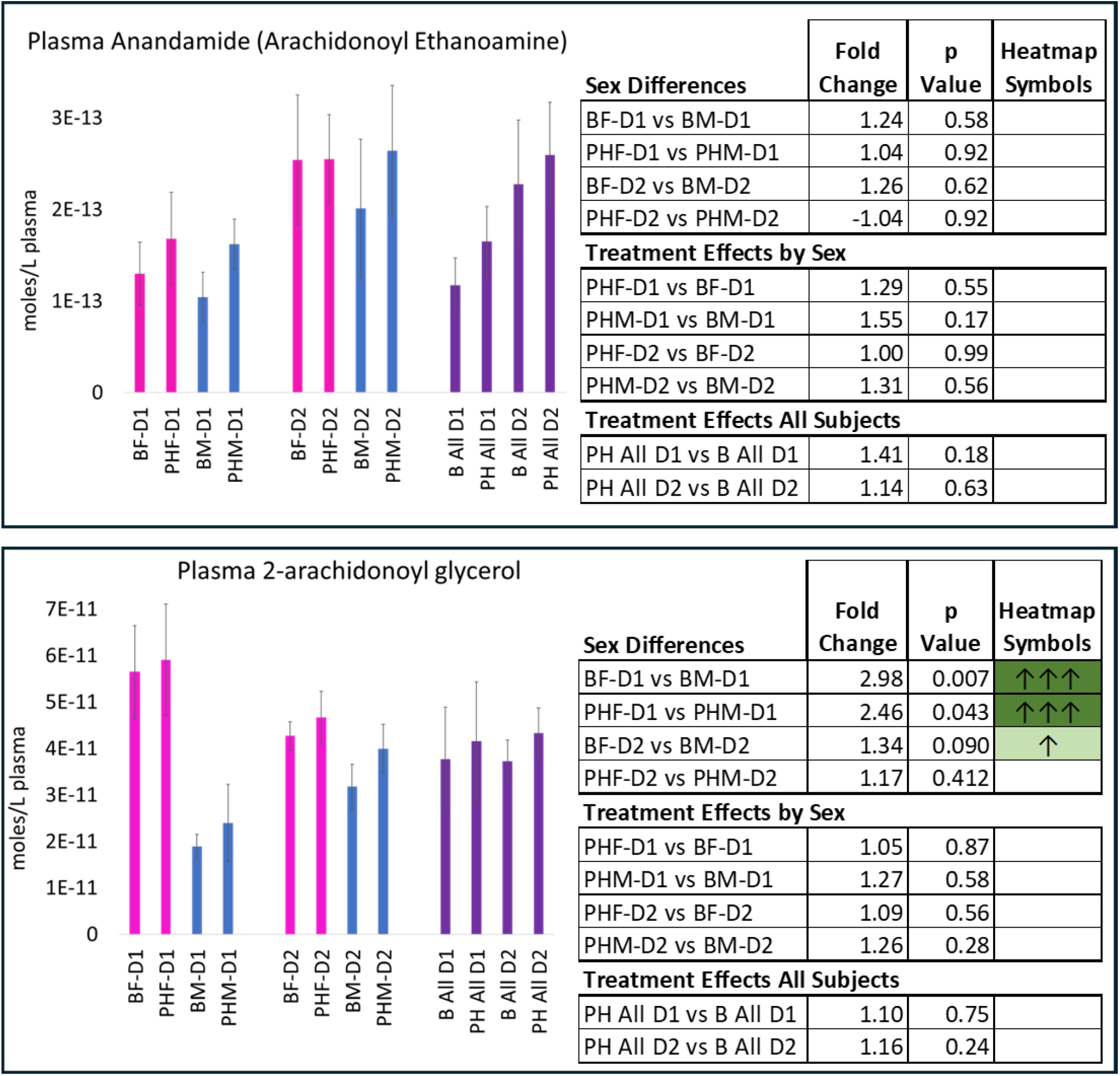
Levels of Anandamide and 2-Arachidonoyl glycerol (2-AG) in plasma at 8 different time points. The bar graphs on the left side are the mean SEM of the levels of endogenous lipids in plasma in each treatment group (Females (F) in pink, Males (M) in blue, and combined (All) in purple). Baseline (B), Post hit (PH), and day 1 and 2 (D1, D2). The table on the right side of the figure lists the analytical data from the interactions of the listed comparisons. *See Methods for description of data analysis*.

**Figure 5.**
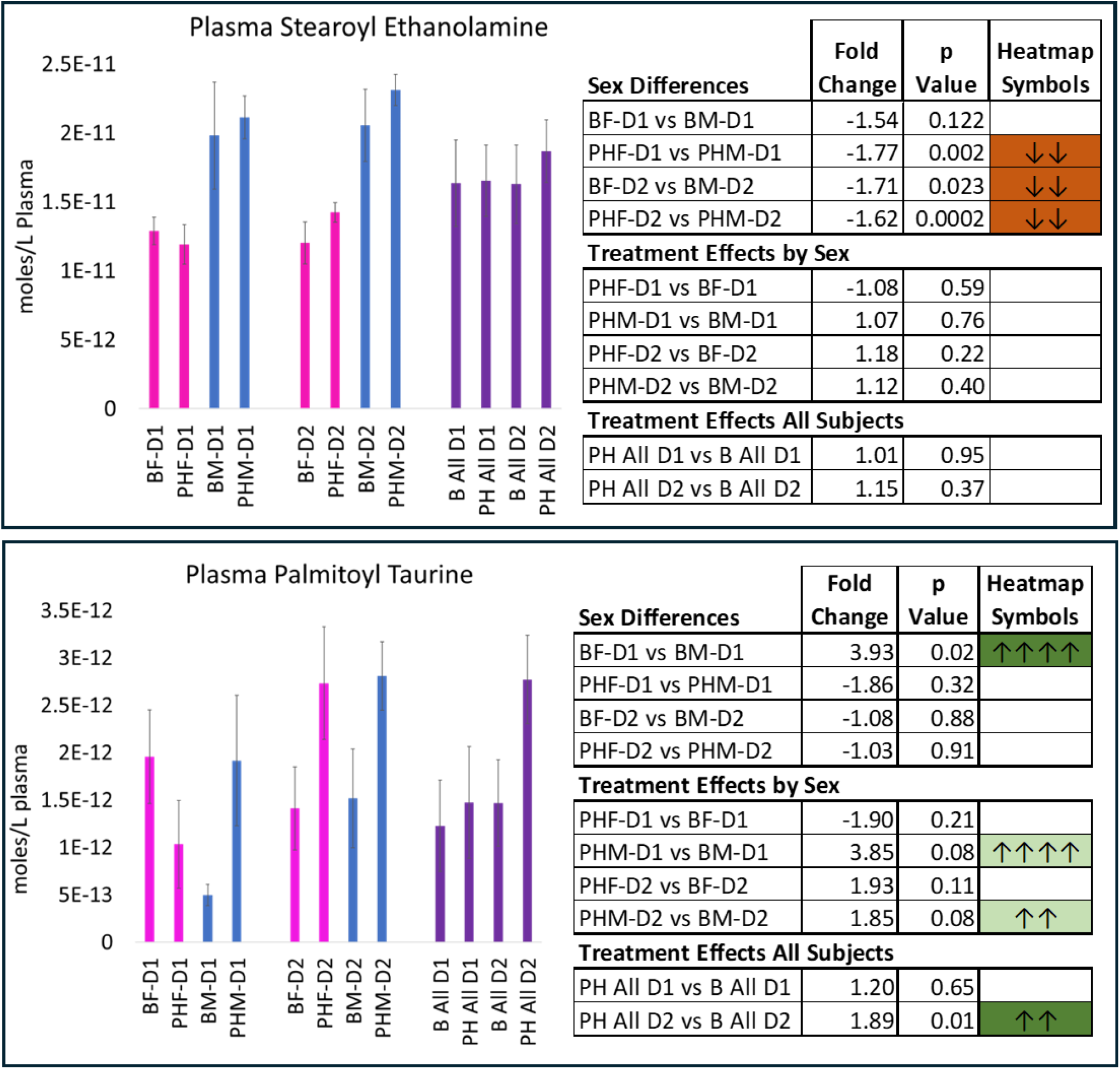
Levels of Stearoyl Ethanolamine and Palmitoyl Taurine in plasma at 8 different time points. The bar graphs on the left side are the mean SEM of the levels of endogenous lipids in plasma in each treatment group (Females (F) in pink, Males (M) in blue, and combined (All) in purple). Baseline (B), Post hit (PH), and day 1 and 2 (D1, D2). The table on the right side of the figure lists the analytical data from the interactions of the listed comparisons. *See Methods for description of data analysis*

**Figure 6.**
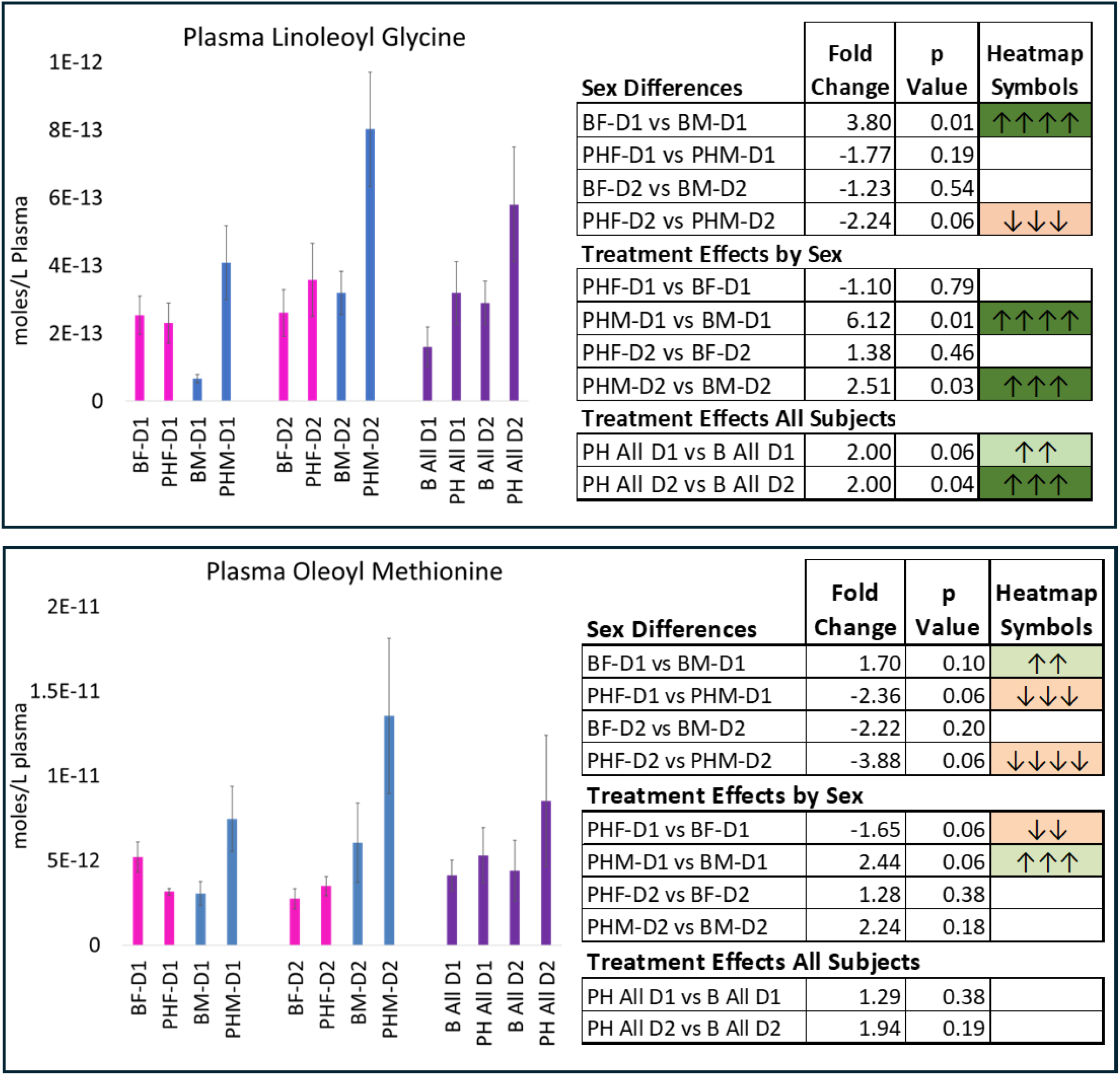
Levels of Linoleoyl Glycine and Oleoyl Methionine in plasma at 8 different time points. The bar graphs on the left side are the mean SEM of the levels of endogenous lipids in plasma in each treatment group (Females (F) in pink, Males (M) in blue, and combined (All) in purple). Baseline (B), Post hit (PH), and day 1 and 2 (D1, D2). The table on the right side of the figure lists the analytical data from the interactions of the listed comparisons. *See Methods for description of data analysis*

Levels of Anandamide (4A) showed no sex differences or treatment differences across within each treatment day. While the bar graphs suggest that levels of AEA increased overall on day 2 of the treatment, we do not have sufficient data to attribute this solely to the mTBI treatment as there is not a day 1 control of no head hit with only restraint and no hit.Therefore, all data are analyzed and reported as the interactions outlined in the tables in Figures 4-6 and are within the day of treatment. Comparisons between treatment days are of the outcomes of these within day analyses. As an example, Figure 4B provides evidence that BF D1 vs. BM D1 and PHF D1 and PHM D1 are significantly different with female 2-AG levels being significantly higher in both conditions. Likewise, BF D2 levels of 2-AG are significantly higher; however, in the p<0.10-0.051 range (p=0.09). Whereas the PHF D2 levels of 2-AG are equivalent to PHM D2 (p=0.412). By contrast, all the within genetic sex comparisons and the combination of females and males of pre-post hit on both D1 and D2 are equivalent.

The close structural congener to Anandamide, stearoyl ethanolamine (SEA), demonstrates a very different expression patterns to either AEA or 2-AG (5A). Opposite to the 2-AG expression, levels of SEA are significantly lower in females than males at 3 of the 4 time points evaluated. However, there was no effect of treatment. With another expression pattern type, palmitoyl taurine (5B) levels are significantly higher in females at baseline on D1 than males but no difference at any other time point. Then, palmitoyl taurine shows significant increases with head hit in males (p=0.08 on both D1 and D2) and females show a trend to increasing after D2 hit (p=0.11), which then translates to a significant increase of p=0.01 on D2 when female and male data are combined.

Plasma levels of linoleoyl glycine (6A) and oleoyl methionine (6B) demonstrate similar patterns to each other; however, different from the first 4 endolipids described here. In both cases, female levels are significantly higher at baseline on D1 to males and then become more equivalent on D2. Likewise, female levels are lower to males after head hit on both days, with fold-change levels being more pronounced and p levels lower for linoleoyl glycine. It is this more pronounced increase in males after head hit that drives the significant increase in linoleoyl glycine in the combined data and not any trending increases by females. That these increases were not as pronounced for oleoyl methionine in males when female levels remained unaffected leads to equivalent levels of oleoyl methionine in the combined data analysis.

We provide these examples to highlight that the heatmap data provides an important snapshot of the overall effects of time, genetic sex, and head hit; however, as with all large-scale data analyses, especially those combining female and male data, should be interpreted withthese types of nuances. Figure 7 provides the heatmap for the endolipids that showed significant changes in at least one analytical interaction. *The heatmap for all lipids screened and the p values for all interactions is in Supplemental Figures*. The data are organized by the acyl chain (*e*.*g*. palmitic acid, linoleic acid, etc.). This overview provides fold-change, significance range, and direction of change for all the lipids summarized in Figure 3. Of note, the direction of difference between females and males reverses after the head hit on day 1; however, the within female analysis shows that no plasma lipids analyzed within 15 minutes of head hit on D1 were significant in the p<0.05 range and this interaction (PH F D1 relative to B F D1) represents comparison with the least overall effect. By dramatic contrast, the effect of head hit on day 1 in males shows the most overall effect of these comparisons and all comparisons show increases in these endolipids. While D2 in males still shows some significant increases there are fewer overall and 2 lipids that decrease. Likewise, D2 in females has an overall expression that resembles the D2 males with mostly increases and 3 decreases.

**Figure 7.**
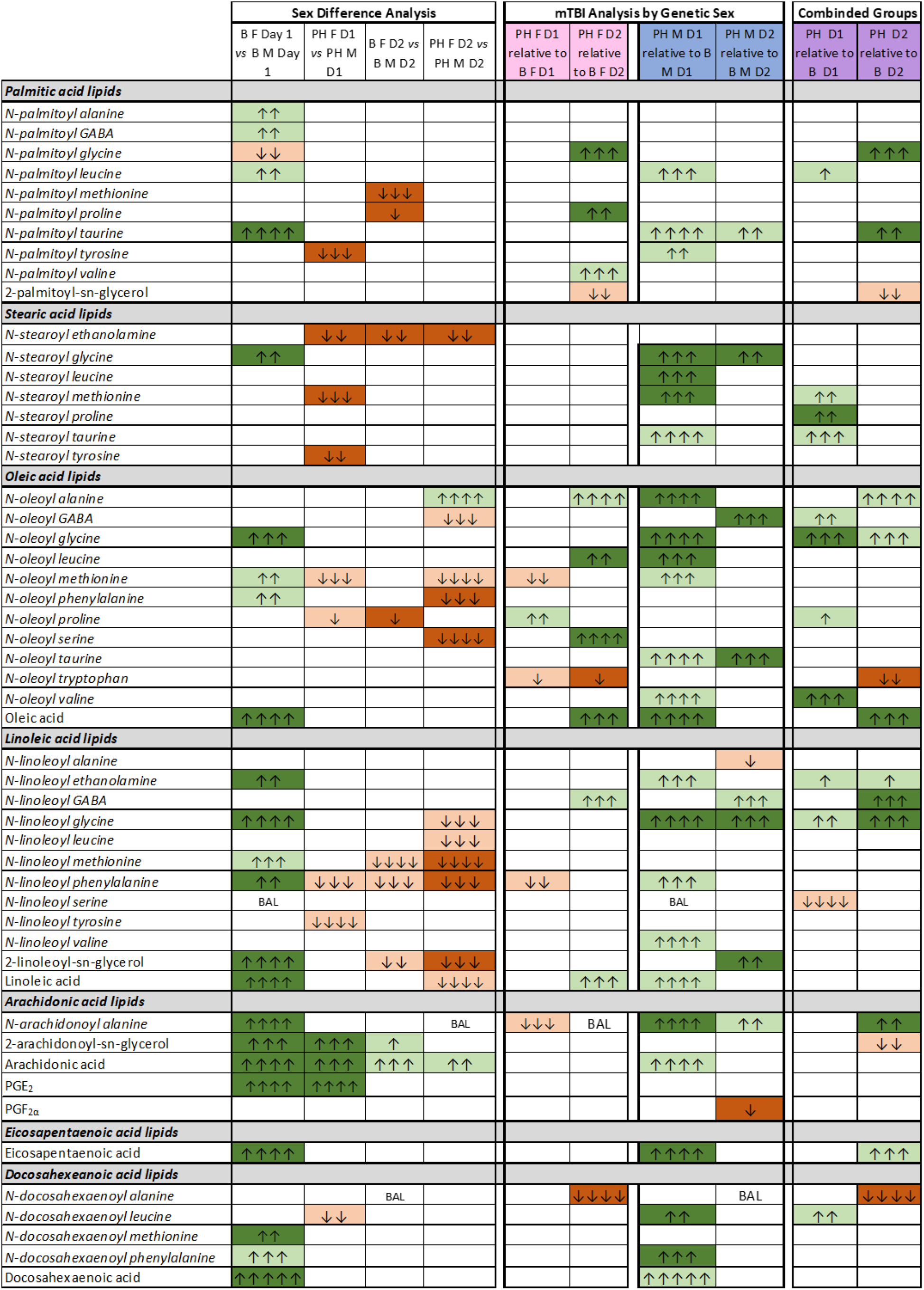
Heatmap of plasma lipids illustrated in 10 different analytical interactions. Data are represented as comparisons of genetic sex at each of the 8 time points (far left columns 1-4); comparisons of pre-post mTBI head hit within genetic sex (columns 5-8) and comparisons of pre-post mTBI head hit with all subjects combined (columns 9-10). Analytical groups are (Females (F), Males (M), Baseline (B), Post hit (PH), and day 1 and 2 (D1, D2). *See Figure 2 for detailed explanation of heatmap fields*. Endogenous lipids are organized by acyl group (fatty acid side chain) and are listed as shortest acyl chain and lowest number of double bonds (palmitic acid C:18:0); to longest acyl chain and most number of double bonds (docosahexaenoic acid C:22:6). Lipoamines/lipo amino acids are designated as the *N*-acyl group amine; then 2-acyl glycerols, and free fatty acids.

## Discussion

These data are an exploratory evaluation of over 80 endolipids, including the eCBs, Anandamide and 2-AG, in plasma directly before and after acute then repeated mTBI injury. These data provide novel data for the potential of evaluating blood biomarkers for predicting mTBI injury effects. Key findings show that there are significant sex differences in both the baseline levels of these endolipids and how these are modified after injury. Importantly, the largest modulation of plasma endolipids measured after mTBI are primarily increases in plasma lipid concentrations and are primarily observed in males.

### Sex differences in effects of mTBI

The notion of biological sex being a modulator of mTBI outcomes is not a new idea. Epidemiological data suggest that females tend to report more negative outcomes over time than males after head trauma; however, most direct comparisons of how males and females differ is after acute insult [1, 38-40]. Bazarian and colleagues suggested that mTBI affects the regulation of hormones, progesterone, and estrogen, as a potential explanation for poorer mTBI outcomes for women [25]. Men do produce these hormones as well, but in much smaller amounts [41], suggesting there may be a production or concentration-dependent effect. In addition to our previous study showing an increase in diffusion coefficients primary in females over time after repeated mTBI [27], Wright and colleagues show that MRI analysis on the subjects after repeated mTBI and found that females experienced greater atrophy of the prefrontal cortex and that there were also differences in the expression of mRNA for various proteins, including tau protein, which is commonly associated with Alzheimer’s Disease [42].

Male subjects can also show deficits after mTBI. Tucker and colleagues investigated the long-term effects of repeated mTBI in mice and showed that impairments remained for up to 6 months after injury, with males showing more significant motor deficits than females [43]. Whereas, the study by Wright and colleagues also conducted cognitive-behavioral tests over time and found that mTBI led to depressive-like behavior in female rats but cognitive deficits in male rats [42]. The effect of aging is also an important factor to consider. A recent study that utilized only male rats found that when compared to older subjects, young mice showed worse physiological outcomes and postulated it may be due to an incomplete development of an immune or inflammatory response [44]. An alternative explanation could be that there was a much more robust immune response due to more collateral damage from microglia [45]. A clinical report somewhat converges on these reports as it suggests that a single mTBI event can begin neurodegeneration that will persist with aging, which can lead to increased vulnerability for mTBI at older ages [46]. Here, we also introduce the idea that the baseline sex differences in lipid signaling molecules may be an additional avenue to explore when investigating how mTBI-inducing injuries can have differential outcomes.

### Changes in eCBs and their congeners with mTBI

Anandamide and 2-AG are the most well-known eCBs and have been cited to have a variety of functions, including neuroprotective roles especially relevant for mTBI. Data in this current study shows that both males and females experienced no overall acute effect from mTBI in plasma eCBs. However, this does not preclude eCBs changing in other parts of the CNS and body. Previous findings from Panikashvili and colleagues have shown 2-AG increases in the brain during periods of injury [47] and in plasma during inflammation [48]. Here, we focused on acute changes in plasma within a time frame of acute injury or re-injury and not longer-term evaluation nor CNS effects as in these previous studies.

### Changes in FFAs with mTBI

In addition to the cognitive changes due to head injury, there are also metabolic changes that have yet to be fully characterized. Specifically, the metabolic changes involving free fatty acids that can also lead to prolonged metabolic dysfunction [49], as many of their oxidative products excluding resolvins [50], are pro-inflammatory [51]. All 5 FFAs measured increased after head hit in males on day one, with no increases measured in females. Importantly, baseline levels of FFAs on day one of the head hits show that females have significantly higher levels of FFAs in all 5 species evaluated. This difference was gone at the pre-hit time point on day 2 for all FFAs except arachidonic acid. Essentially, these data indicate that levels of FFAs were only affected in males after the initial head hit. While this may seem counter-intuitive to a protection mechanism as many FFAs are converted to pro-inflammatory mediators, the FFAs eicosapentaenoic and docosahexaenoic acids can be converted to compounds like resolvins, which may act to protect against inflammatory damage [51, 52]. That the significantly higher levels of FFAs in females at baseline was not present in day 2 baseline could be an effect of the treatment on day one, of overall stress of the procedure, of hormonal cycle differences, or a combination of all of these factors. As an exploratory study, the aim was to first determine if acute and repeated heat hits could alter plasma lipid signaling molecules and if this is sex dependent. Given that the data here clearly illustrate that they do with day 1 analysis alone, the complexity of repeated head hits with potential stress and hormonal modulations of the lipids can be addressed in future studies.

### Changes in lipo amino acids with mTBI

Lipo amino acids refer to the endolipid congeners of *N*-acyl ethanolamines that are conjugations of free fatty acids and amino acids. The data here highlight modulations in multiple conjugation constructs including *N*-acyl glycine, *N*-acyl taurine, *N*-acyl phenylalanine, and *N*-acyl leucine. Prior data from our lab showed that *N*-acyl glycines are upregulated after mTBI in the CNS [21], which agrees with the data in the current study for males (as shown in Figures 6 and 7). Previous data was collected from a male-only cohort so we cannot determine if female CNS levels are likewise affected. Here, data show that the increases in *N*-acyl glycine concentrations after head hit are seen exclusively in males. Some work has implicated the importance of *N*-acyl glycines for neuroprotective effects after traumatic brain injury [45,46] and reperfusion after ischemic stroke [47]. TBI or head injury has the potential to injury vascular networks in the brain [48] and those certain mechanistic responses involving small molecule lipids [49] which offers a potential explanation as to why so many different endolipid species are was so affected due to mTBI.

This pattern of increasing levels of plasma endolipids in males and not females was the same for the data from the *N*-acyl taurines, *N*-acyl leucines, and *N*-acyl methionines. Prior research from our group showed that *N*-arachidonoyl-taurine (A-taur) increased in CNS after mTBI [21], however, plasma levels of A-taur were not in the analytical range in plasma to make this determination. Previous screens were not analyzing additional *N*-acyl taurines in the CNS, so the data here indicating that the more abundant plasma *N*-palmitoyl-taurine and *N*-stearoyl-taurine measured a 4-fold increase in concentration in males after the first day of mTBI indicated that this family of lipids appear to be upregulated after head hit in multiple areas of the body. The literature on the regulation of *N*-acyl taurines is sparse, and not much is known about their signaling roles other than to activate some TRP channels [53], therefore this family of endogenous lipids provides a novel direction for future study.

### Development of lipid biomarkers in plasma as disease outcome predictors

Utilizing a quantifiable, endogenous indicators of an illness or disease has assisted healthcare providers with assessing potential outcomes of their patients’ illnesses, which can range from cancer to neurodegenerative diseases. However, with mTBI, determining these biomarkers and then linking them to future outcomes has been more difficult due to the variable nature of the condition, stemming from the differences in injury and symptomology between individuals. Previous studies have identified potential proteins, enzymes, and amino acids as candidates [6, 54-57], but none to date have proven to be clinically relevant. Other groups have almost exclusively investigated proteins to serve as biomarkers, including the tau protein, glial fibrillary acid protein (GFAP), and the neurofilament protein (NFL), but have also briefly explored enzymes such as hydrolases [54, 57]. Post-injury levels of GFAP and NFL have also been found to be comparable to levels of these two proteins in neurodegenerative brains and have been speculated to accelerate brain aging [55]. Another study has been able to link GFAP and ubiquitin C-terminal hydrolase L1 (UCH-L1) to outcomes of death and other “unfavorable outcomes.” [56]. While these are convincing examples of pre-clinical and clinical models, it is important to note that the main method of obtaining this data included computerized tomography (CT) scans, which involves using low-dose radiation, which may also have an impact on plasma biomarkers, especially if a person receives more than one (*e*.*g*. for multiple injuries). Furthermore, the efficacy of these compounds as biomarkers for mTBI relies on the fluctuations in their levels being noticeably different from other injury types, an aspect that was not addressed in these studies. An examination of a broader range of plasma endolipids as analyzed here could provide additional avenues for evaluations of long-term effects mTBI and related injuries.

Most of endolipids in the screen were present in plasma at a level that provided data for lipidomics analysis using only up to 100µL of plasma, a volume necessary to be able to perform multiple blood draws on these rodent subjects. This threshold provides evidence that the levels of these endolipids could be readily measured continuously throughout a patient’s recovery in that volumes greater than 100µL of plasma can be easily obtained. Therefore, this family of lipids has the potential to provide data on the progression of the injury and could be another crucial aspect in predicting mTBI outcomes.

## Supporting information

Supplemental Heatmaps

## Acknowledgements

**Acknowledgements:** This work was funded by P30DA056410.

## References

1. Alexis B. Peterson, P.M. Karen E. Thomas, and M. Hong Zhou, MPH, Surveillance Report of Traumatic Brain Injury-related Deaths by Age Group, Sex, and Mechanism of Injury—United States, 2018 and 2019. 2022.

2. Cassidy, J.D., et al., Incidence, risk factors and prevention of mild traumatic brain injury: results of the WHO Collaborating Centre Task Force on Mild Traumatic Brain Injury. J Rehabil Med, 2004(43 Suppl): p. 28–60.

3. Agoston, D.V. and M. Elsayed, Serum-based protein biomarkers in blast-induced traumatic brain injury spectrum disorder. Front Neurol, 2012. 3: p. 107.

4. Bahado-Singh, R.O., et al., Serum metabolomic markers for traumatic brain injury: a mouse model. Metabolomics, 2016. 12(6): p. 100.

5. Gill, J., et al., Acute plasma tau relates to prolonged return to play after concussion. Neurology, 2017. 88(6): p. 595–602.

6. Fiandaca, M.S., et al., Plasma metabolomic biomarkers accurately classify acute mild traumatic brain injury from controls. PLoS One, 2018. 13(4): p. e0195318.

7. Kikinis, Z., et al., Diffusion imaging of mild traumatic brain injury in the impact accelerated rodent model: A pilot study. Brain Inj, 2017. 31(10): p. 1376–1381.

8. Oresic, M., et al., Human Serum Metabolites Associate With Severity and Patient Outcomes in Traumatic Brain Injury. EBioMedicine, 2016. 12: p. 118–126.

9. Schurman, L.D. and A.H. Lichtman, Endocannabinoids: A Promising Impact for Traumatic Brain Injury. Front Pharmacol, 2017. 8: p. 69.

10. Shohami, E., et al., Endocannabinoids and traumatic brain injury. Br J Pharmacol, 2011. 163(7): p. 1402–10.

11. Castel, J., et al., NAPE-PLD in the ventral tegmental area regulates reward events, feeding and energy homeostasis. Mol Psychiatry, 2024. 29(5): p. 1478–1490.

12. Warren, G., et al., Discovery and Preclinical Evaluation of a Novel Inhibitor of FABP5, ART26.12, Effective in Oxaliplatin-Induced Peripheral Neuropathy. J Pain, 2024. 25(7): p. 104470.

13. Castel, J., et al., NAPE-PLD in the ventral tegmental area regulates reward events, feeding and energy homeostasis. Res Sq, 2023.

14. Leishman, E., K. Mackie, and H.B. Bradshaw, Elevated Levels of Arachidonic Acid-Derived Lipids Including Prostaglandins and Endocannabinoids Are Present Throughout ABHD12 Knockout Brains: Novel Insights Into the Neurodegenerative Phenotype. Front Mol Neurosci, 2019. 12: p. 142.

15. Leishman, E., et al., Cannabidiol’s Upregulation of N-acyl Ethanolamines in the Central Nervous System Requires N-acyl Phosphatidyl Ethanolamine-Specific Phospholipase D. Cannabis Cannabinoid Res, 2018. 3(1): p. 228–241.

16. Leishman, E., et al., Lipidomics profile of a NAPE-PLD KO mouse provides evidence of a broader role of this enzyme in lipid metabolism in the brain. Biochim Biophys Acta, 2016. 1861(6): p. 491–500.

17. Bradshaw, H.B. and E. Leishman, Lipidomics: A Corrective Lens for Enzyme Myopia. Methods Enzymol, 2017. 593: p. 123–141.

18. Leishman, E., et al., Environmental Toxin Acrolein Alters Levels of Endogenous Lipids, Including TRP Agonists: A Potential Mechanism for Headache Driven by TRPA1 Activation. Neurobiol Pain, 2017. 1: p. 28–36.

19. Johnson, C.T., et al., Cannabinoids accumulate in mouse breast milk and differentially regulate lipid composition and lipid signaling molecules involved in infant development. BBA Adv, 2022. 2.

20. Miller, S., et al., Delta9-Tetrahydrocannabinol and Cannabidiol Differentially Regulate Intraocular Pressure. Invest Ophthalmol Vis Sci, 2018. 59(15): p. 5904–5911.

21. Leishman, E., et al., Bioactive Lipids in Cancer, Inflammation and Related Diseases : Acute and Chronic Mild Traumatic Brain Injury Differentially Changes Levels of Bioactive Lipids in the CNS Associated with Headache. Adv Exp Med Biol, 2019. 1161: p. 193–217.

22. Bashashati, M., et al., Plasma endocannabinoids and cannabimimetic fatty acid derivatives are altered in cyclic vomiting syndrome: The effects of sham feeding. J Investig Med, 2023. 71(8): p. 821–829.

23. Bashashati, M., et al., Plasma endocannabinoids and cannabimimetic fatty acid derivatives are altered in gastroparesis: A sex- and subtype-dependent observation. Neurogastroenterol Motil, 2021. 33(1): p. e13961.

24. Bradshaw, H.B. and C.T. Johnson, Measuring the Content of Endocannabinoid-Like Compounds in Biological Fluids: A Critical Overview of Sample Preparation Methodologies. Methods Mol Biol, 2023. 2576: p. 21–40.

25. Bazarian, J.J., et al., Sex differences in outcome after mild traumatic brain injury. J Neurotrauma, 2010. 27(3): p. 527–39.

26. Aa, D.S., et al., Mild repetitive TBI reduces brain-derived neurotrophic factor (BDNF) in the substantia nigra and hippocampus: A preclinical model for testing BDNF-targeted therapeutics. Exp Neurol, 2024. 374: p. 114696.

27. Bens, N., et al., Multimodal Magnetic Resonance Imaging with Mild Repetitive Head Injury in Awake Rats: Modeling the Human Experience and Clinical Condition. Neurosci Bull, 2025.

28. Kulkarni, P., et al., Neuroradiological Changes Following Single or Repetitive Mild TBI. Front Syst Neurosci, 2019. 13: p. 34.

29. Kilkenny, C., et al., Improving bioscience research reporting: the ARRIVE guidelines for reporting animal research. PLoS Biol, 2010. 8(6): p. e1000412.

30. Viano, D.C., et al., Concussion in professional football: animal model of brain injury--part 15. Neurosurgery, 2009. 64(6): p. 1162-73; discussion 1173.

31. Rowson, S., et al., Biomechanical Perspectives on Concussion in Sport. Sports Med Arthrosc Rev, 2016. 24(3): p. 100–7.

32. Mychasiuk, R., et al., The direction of the acceleration and rotational forces associated with mild traumatic brain injury in rodents effect behavioural and molecular outcomes. J Neurosci Methods, 2016. 257: p. 168–78.

33. Shultz, S.R., et al., Repeated mild lateral fluid percussion brain injury in the rat causes cumulative long-term behavioral impairments, neuroinflammation, and cortical loss in an animal model of repeated concussion. J Neurotrauma, 2012. 29(2): p. 281–94.

34. Xiong, Y., A. Mahmood, and M. Chopp, Animal models of traumatic brain injury. Nat Rev Neurosci, 2013. 14(2): p. 128–42.

35. Aungst, S.L., et al., Repeated mild traumatic brain injury causes chronic neuroinflammation, changes in hippocampal synaptic plasticity, and associated cognitive deficits. J Cereb Blood Flow Metab, 2014. 34(7): p. 1223–32.

36. Fidan, E., et al., Repetitive Mild Traumatic Brain Injury in the Developing Brain: Effects on Long-Term Functional Outcome and Neuropathology. J Neurotrauma, 2016. 33(7): p. 641–51.

37. Stuart, J.M., et al., Brain levels of prostaglandins, endocannabinoids, and related lipids are affected by mating strategies. Int J Endocrinol, 2013. 2013: p. 436252.

38. Albanese, B.J., et al., Anxiety sensitivity mediates gender differences in post-concussive symptoms in a clinical sample. Psychiatry Res, 2017. 252: p. 242–246.

39. Styrke, J., et al., Sex-differences in symptoms, disability, and life satisfaction three years after mild traumatic brain injury: a population-based cohort study. J Rehabil Med, 2013. 45(8): p. 749–57.

40. Mikolic, A., et al., Differences between Men and Women in Treatment and Outcome after Traumatic Brain Injury. J Neurotrauma, 2021. 38(2): p. 235–251.

41. Han, Y., et al., Comparing expression of progesterone and estrogen receptors in testicular tissue from men with obstructive and nonobstructive azoospermia. J Androl, 2009. 30(2): p. 127–33.

42. Wright, D.K., et al., Sex matters: repetitive mild traumatic brain injury in adolescent rats. Ann Clin Transl Neurol, 2017. 4(9): p. 640–654.

43. Tucker, L.B., et al., Chronic Neurobehavioral Sex Differences in a Murine Model of Repetitive Concussive Brain Injury. Front Neurol, 2019. 10: p. 509.

44. Islam, M., et al., Differential neuropathology and functional outcome after equivalent traumatic brain injury in aged versus young adult mice. Exp Neurol, 2021. 341: p. 113714.

45. Chen, Z. and B.D. Trapp, Microglia and neuroprotection. J Neurochem, 2016. 136 Suppl 1: p. 10–7.

46. Tremblay, S., et al., Mild traumatic brain injury: The effect of age at trauma onset on brain structure integrity. Neuroimage Clin, 2019. 23: p. 101907.

47. Panikashvili, D., et al., An endogenous cannabinoid (2-AG) is neuroprotective after brain injury. Nature, 2001. 413(6855): p. 527–31.

48. Panikashvili, D., et al., The endocannabinoid 2-AG protects the blood-brain barrier after closed head injury and inhibits mRNA expression of proinflammatory cytokines. Neurobiol Dis, 2006. 22(2): p. 257–64.

49. Lai, J.Q., et al., Metabolic disorders on cognitive dysfunction after traumatic brain injury. Trends Endocrinol Metab, 2022. 33(7): p. 451–462.

50. Anthonymuthu, T.S., et al., Global assessment of oxidized free fatty acids in brain reveals an enzymatic predominance to oxidative signaling after trauma. Biochim Biophys Acta Mol Basis Dis, 2017. 1863(10 Pt B): p. 2601–2613.

51. Dyall, S.C., Interplay Between n-3 and n-6 Long-Chain Polyunsaturated Fatty Acids and the Endocannabinoid System in Brain Protection and Repair. Lipids, 2017. 52(11): p. 885–900.

52. Serhan, C.N. and N.A. Petasis, Resolvins and protectins in inflammation resolution. Chem Rev, 2011. 111(10): p. 5922–43.

53. Hanus, L., et al., N-Acyl amino acids and their impact on biological processes. Biofactors, 2014. 40(4): p. 381–8.

54. Shahim, P., et al., Neurofilament light and tau as blood biomarkers for sports-related concussion. Neurology, 2018. 90(20): p. e1780–e1788.

55. Newcombe, V.F.J., et al., Post-acute blood biomarkers and disease progression in traumatic brain injury. Brain, 2022. 145(6): p. 2064–2076.

56. Korley, F.K., et al., Prognostic value of day-of-injury plasma GFAP and UCH-L1 concentrations for predicting functional recovery after traumatic brain injury in patients from the US TRACK-TBI cohort: an observational cohort study. Lancet Neurol, 2022. 21(9): p. 803–813.

57. Posti, J.P. and O. Tenovuo, Blood-based biomarkers and traumatic brain injury-A clinical perspective. Acta Neurol Scand, 2022. 146(4): p. 389–399.

