## Supplemental Heatmaps for "Sex Differences in Plasma Levels of Endocannabinoids and Related Lipids Before and After acute and repeated mTBI: an exploratory study for plasma biomarkers for mTBI": 250910_supplemental figures.pdf

| | Sex Difference Analysis | | | | mTBI Analysis by Genetic Sex | | | | Combinded Groups | |
|---|---|---|---|---|---|---|---|---|---|---|
| | B F Day 1 *vs* B M Day 1 | PH F D1 *vs* PH M D1 | B F D2 *vs* B M D2 | PH F D2 *vs* PH M D2 | PH F D1 relative to B F D1 | PH M D1 relative to B M D1 | PH F D2 relative to B F D2 | PH M D2 relative to B M D2 | PH D1 relative to B D1 | PH D2 relative to B D2 |
| ***N-acyl alanine*** | | | | | | | | | | |
| *N-palmitoyl alanine* | 0.0914989 | 0.11338 | 0.8012 | 0.7735923 | 0.266667 | 0.635538 | 0.41224 | 0.804086 | 0.300942 | 0.425334 |
| *N-stearoyl alanine* | 0.8085149 | 0.3835 | 0.2267 | 0.2863002 | 0.77542 | 0.975099 | 0.56357 | 0.895421 | 0.833867 | 0.672447 |
| *N-oleoyl alanine* | 0.1986729 | 0.212 | 0.81444 | 0.0890013 | 0.797045 | 0.006843 | 0.06203 | 0.16544 | 0.447152 | 0.053977 |
| *N-linoleoyl alanine* | 0.2970287 | 0.20297 | 0.67406 | 0.5897751 | 0.364262 | 0.658679 | 0.9407 | 0.098675 | 0.337372 | 0.576987 |
| *N-arachidonoyl alanine* | 0.009553 | 0.11488 | 0.287 | BAL | 0.054027 | 0.037398 | BAL | 0.09496 | 0.988047 | 0.037234 |
| *N-docosahexaenoyl alanine* | 0.4774381 | 0.51034 | BAL | 0.9360986 | 0.853715 | 0.666075 | 0.03221 | BAL | 0.753412 | 0.006771 |
| ***N-acyl ethanolamine*** | | | | | | | | | | |
| *N-palmitoyl ethanolamine* | 0.6709869 | 0.81867 | 0.5763 | 0.6492657 | 0.280359 | 0.49693 | 0.83006 | 0.164523 | 0.202264 | 0.267469 |
| *N-stearoyl ethanolamine* | 0.1219766 | 0.00239 | 0.02265 | 0.0001628 | 0.589955 | 0.759095 | 0.22284 | 0.395816 | 0.951732 | 0.365617 |
| *N-oleoyl ethanolamine* | 0.9864747 | 0.94507 | 0.99823 | 0.6685717 | 0.791963 | 0.688914 | 0.61178 | 0.365937 | 0.617946 | 0.275936 |
| *N-linoleoyl ethanolamine* | 0.0205018 | 0.57082 | 0.8631 | 0.8142833 | 0.844235 | 0.051669 | 0.1847 | 0.337583 | 0.077857 | 0.081553 |
| *N-arachidonoyl ethanolamine* | 0.5752769 | 0.91817 | 0.62387 | 0.915751 | 0.548819 | 0.17171 | 0.99464 | 0.561924 | 0.179155 | 0.627138 |
| *N-docosahexaenoyl ethanolamine* | 0.57508 | 0.17066 | 0.73891 | 0.2300159 | 0.113727 | 0.72248 | 0.10053 | 0.9264 | 0.314844 | 0.189269 |
| ***N-acyl GABA*** | | | | | | | | | | |
| *N-palmitoyl GABA* | 0.0855428 | 0.24004 | 0.85075 | 0.5157836 | 0.432208 | 0.433868 | 0.52518 | 0.784558 | 0.304282 | 0.544328 |
| *N-stearoyl GABA* | 0.2150089 | 0.91229 | 0.67248 | 0.8098876 | 0.162872 | 0.852409 | 0.98409 | 0.851263 | 0.347284 | 0.857218 |
| *N-oleoyl GABA* | 0.8722555 | 0.88669 | 0.48443 | 0.0847902 | 0.107177 | 0.366732 | 0.794 | 0.042109 | 0.087212 | 0.132448 |
| *N-linoleoyl GABA* | 0.9277775 | 0.10498 | 0.62386 | 0.2766856 | 0.822878 | 0.320367 | 0.05774 | 0.076148 | 0.299408 | 0.016863 |
| *N-arachidonoyl GABA* | 0.7404154 | 0.62245 | 0.92814 | 0.6340548 | 0.848857 | 0.468539 | 0.30429 | 0.391196 | 0.412893 | 0.144419 |
| *N-docosahexaenoyl GABA* | 0.8785978 | 0.84818 | 0.91664 | 0.2201941 | 0.977074 | 0.946626 | 0.79502 | 0.255117 | 0.976042 | 0.443387 |
| ***N-acyl glycine*** | | | | | | | | | | |
| *N-palmitoyl glycine* | 0.0591935 | 0.87208 | 0.84912 | 0.9916782 | 0.161391 | 0.59942 | 0.04938 | 0.170831 | 0.151889 | 0.016346 |
| *N-stearoyl glycine* | 0.0411894 | 0.67171 | 0.22743 | 0.2067055 | 0.913289 | 0.025026 | 0.80218 | 0.032814 | 0.209226 | 0.165795 |
| *N-oleoyl glycine* | 0.0131839 | 0.25137 | 0.47028 | 0.229919 | 0.647748 | 0.049095 | 0.31477 | 0.129709 | 0.042055 | 0.067814 |
| *N-linoleoyl glycine* | 0.0116626 | 0.19117 | 0.54108 | 0.0570705 | 0.786058 | 0.014634 | 0.4647 | 0.027827 | 0.055027 | 0.035942 |
| *N-arachidonoyl glycine* | 0.2880452 | BAL | 0.86449 | 0.2767009 | 0.455076 | BAL | 0.77373 | 0.546077 | 0.190049 | 0.935043 |
| *N-docosahexaenoyl glycine* | 0.4762518 | 0.20439 | 0.17806 | 0.494413 | 0.475656 | 0.853542 | 0.35237 | 0.815988 | 0.533314 | 0.403885 |
| ***N-acyl leucine*** | | | | | | | | | | |
| *N-palmitoyl leucine* | 0.0649264 | 0.40396 | 0.46855 | 0.722344 | 0.963387 | 0.057678 | 0.11594 | 0.411742 | 0.090061 | 0.112051 |
| *N-stearoyl leucine* | 0.1050655 | 0.11943 | 0.28744 | 0.2240368 | 0.385889 | 0.036617 | 0.23141 | 0.24662 | 0.244737 | 0.115952 |
| *N-oleoyl leucine* | 0.4260471 | 0.14288 | 0.46839 | 0.5511642 | 0.956173 | 0.031391 | 0.03341 | 0.4195 | 0.123803 | 0.134539 |
| *N-linoleoyl leucine* | 0.7266319 | 0.19905 | 0.11439 | 0.060133 | 0.300876 | 0.366487 | 0.24337 | 0.289814 | 0.825959 | 0.244366 |
| *N-docosahexaenoyl leucine* | 0.9352569 | 0.06676 | 0.63002 | 0.3536035 | 0.757157 | 0.029862 | 0.54263 | 0.287617 | 0.091875 | 0.93413 |
| ***N-acyl methionine*** | | | | | | | | | | |
| *N-palmitoyl methionine* | 0.8227732 | 0.25523 | 0.02878 | 0.2339802 | 0.985681 | 0.196511 | 0.18127 | 0.743512 | 0.255927 | 0.401475 |
| *N-stearoyl methionine* | 0.1010347 | 0.0219 | 0.2453 | 0.8244894 | 0.43248 | 0.009103 | 0.27976 | 0.44556 | 0.086761 | 0.165839 |
| *N-oleoyl methionine* | 0.0959928 | 0.0563 | 0.20444 | 0.0615256 | 0.055845 | 0.063238 | 0.37905 | 0.183884 | 0.38348 | 0.189604 |
| *N-linoleoyl methionine* | 0.0519419 | 0.24764 | 0.09002 | 0.0097455 | 0.401557 | 0.124768 | 0.98877 | 0.133943 | 0.211558 | 0.24294 |
| *N-arachidonoyl methionine* | 0.975967 | 0.79425 | 0.2122 | 0.6138283 | 0.763083 | 0.953526 | 0.67028 | 0.379768 | 0.753345 | 0.381688 |
| *N-docosahexaenoyl methionine* | 0.010272 | 0.92373 | 0.2006 | 0.3477691 | 0.529653 | 0.192452 | 0.34367 | 0.512812 | 0.948032 | 0.261817 |
| **N- acyl phenylalanine** | | | | | | | | | | |
| *N-palmitoyl phenylalanine* | 0.1767055 | 0.56366 | 0.62981 | 0.2200342 | 0.868 | 0.148925 | 0.27745 | 0.225765 | 0.16727 | 0.105411 |
| *N-stearoyl phenylalanine* | 0.9528506 | 0.17199 | 0.6447 | 0.2627822 | 0.684856 | 0.199814 | 0.77446 | 0.478436 | 0.337448 | 0.431433 |
| *N-oleoyl phenylalanine* | 0.0921463 | 0.23549 | 0.47708 | 0.0389219 | 0.184134 | 0.124954 | 0.77562 | 0.17255 | 0.755124 | 0.199291 |
| *N-linoleoyl phenylalanine* | 0.0488714 | 0.08905 | 0.07485 | 0.0221544 | 0.06337 | 0.071075 | 0.89088 | 0.452513 | 0.299115 | 0.589137 |
| *N-arachidonoyl phenylalanine* | 0.2907379 | 0.87382 | 0.39004 | 0.2359235 | 0.322738 | 0.807412 | 0.49928 | 0.718806 | 0.412574 | 0.513012 |
| *N-docosahexaenoyl phenylalanine* | 0.0574874 | 0.1459 | 0.29908 | 0.7629228 | 0.198821 | 0.04168 | 0.67375 | 0.805879 | 0.537375 | 0.793361 |

P=values of all interactions listed.

| | Sex Difference Analysis | | | | mTBI Analysis by Genetic Sex | | | | Combinded Groups | |
|---|---|---|---|---|---|---|---|---|---|---|
| | B F Day 1 *vs* B M Day 1 | PH F D1 *vs* PH M D1 | B F D2 *vs* B M D2 | PH F D2 *vs* PH M D2 | PH F D1 relative to B F D1 | PH M D1 relative to B M D1 | PH F D2 relative to B F D2 | PH M D2 relative to B M D2 | PH D1 relative to B D1 | PH D2 relative to B D2 |
| ***N*- acyl proline** | | | | | | | | | | |
| *N-palmitoyl proline* | 0.80348 | 0.599095 | 0.023191 | 0.405005 | 0.44687 | 0.109299 | 0.032497 | 0.89413 | 0.100727 | 0.11993 |
| *N-stearoyl proline* | BAL | BAL | BAL | BAL | BAL | BAL | BAL | BAL | 0.014864 | 0.396919 |
| *N-oleoyl proline* | 0.182925 | 0.0676 | 0.023787 | 0.81131 | 0.062058 | 0.171742 | 0.554169 | 0.250353 | 0.050241 | 0.659085 |
| *N-linoleoyl proline* | 0.177759 | 0.284585 | 0.491309 | 0.766931 | 0.172164 | 0.366521 | 0.282828 | 0.37928 | 0.640456 | 0.200887 |
| *N-arachidonoyl proline* | 0.82596 | 0.893055 | 0.014274 | 0.549048 | 0.918244 | 0.727803 | 0.438488 | 0.500007 | 0.752327 | 0.694637 |
| *N-docosahexaenoyl proline* | 0.526658 | 0.922449 | 0.576514 | 0.537944 | 0.658352 | 0.640282 | 0.971808 | 0.868348 | 0.812524 | 0.785997 |
| ***N-acyl serine*** | | | | | | | | | | |
| *N-palmitoyl serine* | 0.309287 | 0.465478 | 0.218613 | 0.19618 | 0.380434 | 0.253589 | 0.641952 | 0.597976 | 0.234378 | 0.746309 |
| *N-stearoyl serine* | 0.831469 | 0.433992 | 0.934762 | 0.394288 | 0.777495 | 0.473277 | 0.356261 | 0.928052 | 0.701435 | 0.384456 |
| *N-oleoyl serine* | 0.151832 | 0.305855 | 0.367703 | 0.033514 | 0.634145 | 0.181561 | 0.04588 | 0.553294 | 0.15868 | 0.575248 |
| *N-linoleoyl serine* | BAL | 0.783349 | 0.35143 | 0.224011 | 0.110418 | BAL | 0.340052 | 0.71465 | 0.099657 | 0.472325 |
| *N-arachidonoyl serine* | BAL | BAL | BAL | BAL | BAL | BAL | BAL | BAL | 0.598752 | 0.475567 |
| *N-docosahexaenoyl serine* | BAL | BAL | 0.887362 | BAL | BAL | BAL | BAL | 0.949191 | 0.179906 | 0.277093 |
| ***N-acyl taurine*** | | | | | | | | | | |
| *N-palmitoyl taurine* | 0.020723 | 0.318364 | 0.880112 | 0.914847 | 0.20875 | 0.077078 | 0.1123 | 0.076462 | 0.653982 | 0.010934 |
| *N-stearoyl taurine* | 0.230627 | 0.203998 | 0.206638 | 0.806366 | 0.670596 | 0.051201 | 0.479752 | 0.322056 | 0.054048 | 0.403719 |
| *N-oleoyl taurine* | 0.398912 | 0.607436 | 0.891555 | 0.168045 | 0.63696 | 0.09711 | 0.748573 | 0.026466 | 0.115407 | 0.113573 |
| *N-arachidonoyl taurine* | 0.371115 | 0.302235 | 0.415131 | 0.938669 | 0.273406 | 0.388786 | 0.761522 | 0.516088 | 0.712361 | 0.679061 |
| ***N-acyl tryptophan*** | | | | | | | | | | |
| *N-palmitoyl tryptophan* | 0.39215 | 0.739492 | 0.746118 | 0.4225 | 0.898403 | 0.414397 | 0.878526 | 0.5685 | 0.49956 | 0.508918 |
| *N-stearoyl tryptophan* | 0.518025 | 0.604261 | 0.889812 | 0.620532 | 0.360812 | 0.78383 | 0.905116 | 0.618784 | 0.625332 | 0.751559 |
| *N-oleoyl tryptophan* | 0.888869 | 0.232148 | 0.305089 | 0.724921 | 0.097716 | 0.408913 | 0.026637 | 0.163666 | 0.427012 | 0.043395 |
| *N-linoleoyl tryptophan* | BAL | BAL | BAL | 0.758078 | BAL | BAL | BAL | 0.445158 | 0.267099 | 0.517199 |
| *N-arachidonoyl tryptophan* | BAL | BAL | BAL | BAL | BAL | BAL | BAL | BAL | BAL | BAL |
| *N-docosahexaenoyl tryptophan* | BAL | BAL | BAL | BAL | BAL | BAL | BAL | BAL | BAL | BAL |
| ***N-acyl tyrosine*** | | | | | | | | | | |
| *N-palmitoyl tyrosine* | 0.399811 | 0.035359 | 0.185107 | 0.19126 | 0.219295 | 0.075844 | 0.590724 | 0.481165 | 0.809043 | 0.395883 |
| *N-stearoyl tyrosine* | 0.932152 | 0.011422 | 0.115657 | 0.383099 | 0.322523 | 0.520417 | 0.542888 | 0.853737 | 0.730047 | 0.699094 |
| *N-oleoyl tyrosine* | 0.317214 | 0.185047 | 0.142739 | 0.100529 | 0.155412 | 0.2885 | 0.429623 | 0.369175 | 0.90973 | 0.308727 |
| *N-linoleoyl tyrosine* | 0.856194 | 0.050422 | 0.288653 | 0.109218 | 0.233188 | 0.298736 | 0.468057 | 0.401588 | 0.863565 | 0.578065 |
| *N-arachidonoyl tyrosine* | 0.554551 | 0.106136 | 0.414005 | 0.616466 | 0.435368 | 0.861721 | 0.42204 | 0.4447 | 0.432251 | 0.361563 |
| *N-docosahexaenoyl tyrosine* | 0.914238 | BAL | BAL | 0.536073 | BAL | 0.762824 | BAL | 0.602439 | 0.757579 | 0.737982 |
| ***N-acyl valine*** | | | | | | | | | | |
| *N-palmitoyl valine* | 0.541202 | 0.361286 | 0.152726 | 0.453991 | 0.868513 | 0.161056 | 0.059554 | 0.618637 | 0.185823 | 0.234406 |
| *N- stearoyl valine* | 0.944498 | 0.982431 | 0.308585 | 0.870956 | 0.156125 | 0.380906 | 0.169427 | 0.531923 | 0.112959 | 0.123495 |
| *N-oleoyl valine* | 0.762988 | 0.47681 | 0.211911 | 0.268612 | 0.238265 | 0.063272 | 0.160392 | 0.518424 | 0.02064 | 0.268957 |
| *N-linoleoyl valine* | 0.103169 | 0.237048 | 0.179819 | 0.162777 | 0.536002 | 0.090206 | 0.749571 | 0.293107 | 0.213514 | 0.300669 |
| *N-docosahexaenoyl valine* | 0.758426 | 0.265557 | 0.768434 | 0.923247 | 0.832905 | 0.242907 | 0.899128 | 0.764858 | 0.303743 | 0.980941 |
| **2-acyl-sn-glycerol** | | | | | | | | | | |
| 2-palmitoyl-sn-glycerol | 0.997327 | 0.112857 | 0.882771 | 0.518711 | 0.116983 | 0.94334 | 0.056548 | 0.518999 | 0.200301 | 0.090133 |
| 2-oleoyl-sn-glycerol | 0.219627 | 0.488621 | 0.346903 | 0.357726 | 0.494455 | 0.424137 | 0.50839 | 0.180704 | 0.709664 | 0.132118 |
| 2-linoleoyl-sn-glycerol | 0.019013 | 0.628299 | 0.068801 | 0.037569 | 0.513982 | 0.193971 | 0.81928 | 0.043222 | 0.647235 | 0.148422 |
| 2-arachidonoyl-sn-glycerol | 0.006898 | 0.869403 | 0.576454 | 0.042974 | 0.090298 | 0.555761 | 0.281228 | 0.412267 | 0.751932 | 0.239502 |
| **Free Fatty Acids** | | | | | | | | | | |
| Oleic acid | 0.030587 | 0.776932 | 0.591931 | 0.212559 | 0.875923 | 0.026508 | 0.046254 | 0.123226 | 0.168675 | 0.040026 |
| Linoleic acid | 0.005684 | 0.848046 | 0.318277 | 0.089935 | 0.490937 | 0.058091 | 0.097065 | 0.133351 | 0.341241 | 0.107855 |
| Arachidonic acid | 0.01566 | 0.043004 | 0.066499 | 0.070938 | 0.301848 | 0.054579 | 0.138313 | 0.367853 | 0.797528 | 0.137097 |
| Eicosapentaenoic acid | 0.023656 | 0.132021 | 0.870971 | 0.27117 | 0.238033 | 0.02033 | 0.330742 | 0.123526 | 0.305594 | 0.060198 |
| Docosahexaenoic acid | 0.037334 | 0.636596 | 0.343913 | 0.258797 | 0.145018 | 0.088146 | 0.235119 | 0.249597 | 0.658521 | 0.115852 |
| **Prostaglandins** | | | | | | | | | | |
| $PGE_2$ | 0.024026 | 0.036419 | 0.173964 | 0.297523 | 0.950321 | 0.843496 | 0.939287 | 0.729764 | 0.611016 | 0.779337 |
| $PGF_{2\alpha}$ | 0.213595 | 0.118718 | 0.385731 | 0.109905 | 0.375166 | 0.258955 | 0.64172 | 0.019181 | 0.15962 | 0.348514 |
| 6-keto-$PGF_{1\alpha}$ | BAL | BAL | BAL | BAL | BAL | BAL | BAL | BAL | BAL | BAL |

| | B F Day 1 *vs* B M Day 1 | PH F D1 *vs* PH M D1 | B F D2 *vs* B M D2 | PH F D2 *vs* PH M D2 | PH F D1 relative to B F D1 | PH M D1 relative to B M D1 | PH F D2 relative to B F D2 | PH M D2 relative to B M D2 | PH D1 relative to B D1 | PH D2 relative to B D2 |
|---|---|---|---|---|---|---|---|---|---|---|
| ***N-acyl alanine*** | | | | | | | | | | |
| *N-palmitoyl alanine* | ↑↑ | | | | | | | | | |
| *N-stearoyl alanine* | | | | | | | | | | |
| *N-oleoyl alanine* | | | | ↑↑↑↑ | | ↑↑↑↑ | ↑↑↑↑ | | | ↑↑↑↑ |
| *N-linoleoyl alanine* | | | | | | | | ↓ | | |
| *N-arachidonoyl alanine* | ↑↑↑↑ | | | BAL | ↓↓↓ | ↑↑↑↑ | BAL | ↑↑ | | ↑↑ |
| *N-docosahexaenoyl alanine* | | | BAL | | | | ↓↓↓↓ | BAL | | ↓↓↓↓ |
| ***N-acyl ethanolamine*** | | | | | | | | | | |
| *N-palmitoyl ethanolamine* | | | | | | | | | | |
| *N-stearoyl ethanolamine* | | ↓↓ | ↓↓ | ↓↓ | | | | | | |
| *N-oleoyl ethanolamine* | | | | | | | | | | |
| *N-linoleoyl ethanolamine* | ↑↑ | | | | | ↑↑↑ | | | ↑ | ↑ |
| *N-arachidonoyl ethanolamine* | | | | | | | | | | |
| *N-docosahexaenoyl ethanolamine* | | | | | | | | | | |
| ***N-acyl GABA*** | | | | | | | | | | |
| *N-palmitoyl GABA* | ↑↑ | | | | | | | | | |
| *N-stearoyl GABA* | | | | | | | | | | |
| *N-oleoyl GABA* | | | | ↓↓↓ | | | | ↑↑↑ | ↑↑ | |
| *N-linoleoyl GABA* | | | | | | | ↑↑↑ | ↑↑↑ | | ↑↑↑ |
| *N-arachidonoyl GABA* | | | | | | | | | | |
| *N-docosahexaenoyl GABA* | | | | | | | | | | |
| ***N-acyl glycine*** | | | | | | | | | | |
| *N-palmitoyl glycine* | ↓↓ | | | | | | ↑↑↑ | | | ↑↑↑ |
| *N-stearoyl glycine* | ↑↑ | | | | | ↑↑↑ | | ↑↑ | | |
| *N-oleoyl glycine* | ↑↑↑ | | | | | ↑↑↑↑ | | | ↑↑↑ | ↑↑↑ |
| *N-linoleoyl glycine* | ↑↑↑↑ | | | ↓↓↓ | | ↑↑↑↑ | | ↑↑↑ | ↑↑ | ↑↑↑ |
| *N-arachidonoyl glycine* | | BAL | | | | BAL | | | | |
| *N-docosahexaenoyl glycine* | | | | | | | | | | |
| ***N-acyl leucine*** | | | | | | | | | | |
| *N-palmitoyl leucine* | ↑↑ | | | | | ↑↑↑ | | | ↑ | |
| *N-stearoyl leucine* | | | | | | ↑↑↑ | | | | |
| *N-oleoyl leucine* | | | | | | ↑↑↑ | ↑↑ | | | |
| *N-linoleoyl leucine* | | | | ↓↓↓ | | | | | | |
| *N-docosahexaenoyl leucine* | | ↓↓ | | | | ↑↑ | | | ↑↑ | |
| ***N-acyl methionine*** | | | | | | | | | | |
| *N-palmitoyl methionine* | | | ↓↓↓ | | | | | | | |
| *N-stearoyl methionine* | | ↓↓↓ | | | | ↑↑↑ | | | ↑↑ | |
| *N-oleoyl methionine* | ↑↑ | ↓↓↓ | | ↓↓↓↓ | ↓↓ | ↑↑↑ | | | | |
| *N-linoleoyl methionine* | ↑↑↑ | | ↓↓↓↓ | ↓↓↓↓ | | | | | | |
| *N-arachidonoyl methionine* | | | | | | | | | | |
| *N-docosahexaenoyl methionine* | ↑↑ | | | | | | | | | |
| ***N- acyl phenylalanine*** | | | | | | | | | | |
| *N-palmitoyl phenylalanine* | | | | | | | | | | |
| *N-stearoyl phenylalanine* | | | | | | | | | | |
| *N-oleoyl phenylalanine* | ↑↑ | | | ↓↓↓ | | | | | | |
| *N-linoleoyl phenylalanine* | ↑↑ | ↓↓↓ | ↓↓↓ | ↓↓↓ | ↓↓ | ↑↑↑ | | | | |
| *N-arachidonoyl phenylalanine* | | | | | | | | | | |
| *N-docosahexaenoyl phenylalanine* | ↑↑↑ | | | | | ↑↑↑ | | | | |

Heatmaps of all interactions listed.

| | Sex Difference Analysis | | | | mTBI Analysis by Genetic Sex | | | | Combinded Groups | |
|---|---|---|---|---|---|---|---|---|---|---|
| | B F Day 1 *vs* B M Day 1 | PH F D1 *vs* PH M D1 | B F D2 *vs* B M D2 | PH F D2 *vs* PH M D2 | PH F D1 relative to B F D1 | PH M D1 relative to B M D1 | PH F D2 relative to B F D2 | PH M D2 relative to B M D2 | PH D1 relative to B D1 | PH D2 relative to B D2 |
| ***N*-acyl proline** | | | | | | | | | | |
| *N-palmitoyl proline* | | | ↓ | | | | ↑↑ | | | |
| *N-stearoyl proline* | BAL | BAL | BAL | BAL | BAL | BAL | BAL | BAL | ↑↑ | |
| *N-oleoyl proline* | | ↓ | ↓ | | ↑↑ | | | | ↑ | |
| *N-linoleoyl proline* | | | | | | | | | | |
| *N-arachidonoyl proline* | | | | | | | | | | |
| *N-docosahexaenoyl proline* | | | | | | | | | | |
| ***N-acyl serine*** | | | | | | | | | | |
| *N-palmitoyl serine* | | | | | | | | | | |
| *N-stearoyl serine* | | | | | | | | | | |
| *N-oleoyl serine* | | | | ↓↓↓↓ | | | ↑↑↑↑ | | | |
| *N-linoleoyl serine* | BAL | | | | | BAL | | | ↓↓↓↓ | |
| *N-arachidonoyl serine* | BAL | BAL | BAL | BAL | BAL | BAL | BAL | BAL | | |
| *N-docosahexaenoyl serine* | BAL | BAL | | BAL | BAL | BAL | BAL | | | |
| ***N-acyl taurine*** | | | | | | | | | | |
| *N-palmitoyl taurine* | ↑↑↑↑ | | | | | ↑↑↑↑ | | ↑↑ | | ↑↑ |
| *N-stearoyl taurine* | | | | | | ↑↑↑↑ | | | ↑↑↑ | |
| *N-oleoyl taurine* | | | | | | ↑↑↑↑ | | ↑↑↑ | | |
| *N-arachidonoyl taurine* | | | | | | | | | | |
| ***N-acyl tryptophan*** | | | | | | | | | | |
| *N-palmitoyl tryptophan* | | | | | | | | | | |
| *N-stearoyl tryptophan* | | | | | | | | | | |
| *N-oleoyl tryptophan* | | | | | ↓ | | ↓ | | | ↓↓ |
| *N-linoleoyl tryptophan* | BAL | BAL | BAL | | BAL | BAL | BAL | | | |
| *N-arachidonoyl tryptophan* | BAL | BAL | BAL | BAL | BAL | BAL | BAL | BAL | BAL | BAL |
| *N-docosahexaenoyl tryptophan* | BAL | BAL | BAL | BAL | BAL | BAL | BAL | BAL | BAL | BAL |
| ***N-acyl tyrosine*** | | | | | | | | | | |
| *N-palmitoyl tyrosine* | | ↓↓↓ | | | | ↑↑ | | | | |
| *N-stearoyl tyrosine* | | ↓↓ | | | | | | | | |
| *N-oleoyl tyrosine* | | | | | | | | | | |
| *N-linoleoyl tyrosine* | | ↓↓↓↓ | | | | | | | | |
| *N-arachidonoyl tyrosine* | | | | | | | | | | |
| *N-docosahexaenoyl tyrosine* | | BAL | BAL | | BAL | | BAL | | | |
| ***N-acyl valine*** | | | | | | | | | | |
| *N-palmitoyl valine* | | | | | | | ↑↑↑ | | | |
| *N- stearoyl valine* | | | | | | | | | | |
| *N-oleoyl valine* | | | | | | ↑↑↑↑ | | | ↑↑↑ | |
| *N-linoleoyl valine* | | | | | | ↑↑↑↑ | | | | |
| *N-docosahexaenoyl valine* | | | | | | | | | | |
| **2-acyl-sn-glycerol** | | | | | | | | | | |
| 2-arachidonoyl-sn-glycerol | | | | | | | ↓↓ | | | ↓↓ |
| 2-palmitoyl-sn-glycerol | | | | | | | | | | |
| 2-oleoyl-sn-glycerol | ↑↑↑↑ | | ↓↓ | ↓↓↓ | | | | ↑↑ | | |
| 2-linoleoyl-sn-glycerol | ↑↑↑ | | | ↑↑↑ | ↑ | | | | | |
| **Free Fatty Acids** | | | | | | | | | | |
| Oleic acid | ↑↑↑↑ | | | | | ↑↑↑↑ | ↑↑↑ | | | ↑↑↑ |
| Linoleic acid | ↑↑↑↑ | | | ↓↓↓↓ | | ↑↑↑↑ | ↑↑↑ | | | |
| Arachidonic acid | ↑↑↑↑ | ↑↑↑ | ↑↑↑ | ↑↑ | | ↑↑↑↑ | | | | |
| Eicosapentaenoic acid | ↑↑↑↑ | | | | | ↑↑↑↑ | | | | ↑↑↑ |
| Docosahexaenoic acid | ↑↑↑↑↑ | | | | | ↑↑↑↑↑ | | | | |
| **Prostaglandins** | | | | | | | | | | |
| $PGE_2$ | ↑↑↑↑ | ↑↑↑↑ | | | | | | | | |
| $PGF_{2\alpha}$ | | | | | | | | ↓ | | |
| 6-keto-$PGF_{1\alpha}$ | BAL | BAL | BAL | BAL | BAL | BAL | BAL | BAL | BAL | BAL |
